# A mechanism for synaptic copy between neural circuits

**DOI:** 10.1101/351114

**Authors:** Yuxiu Shao, Binxu Wang, Andrew T. Sornborger, Louis Tao

## Abstract

The brain has a central, short-term learning module, the hippocampus, which transfers what it has learned to long-term memory in cortex during non-REM sleep. The putative mechanism responsible for this type of memory consolidation invokes hierarchically nested hippocampal ripples (100-250 Hz), thalamo-cortical spindles (7-15 Hz), and cortical slow oscillations (< 1 Hz) to enable transfer. Suppression of, for instance, thalamic spindles has been shown to impair hippocampus-dependent memory consolidation. Cortical oscillations are central to information transfer in neural systems. Significant evidence supports the idea that coincident spike input can allow the neural threshold to be overcome, and spikes to be propagated downstream in a circuit. Thus, an observation of oscillations in neural circuits would be an indication that repeated synchronous spiking is enabling information transfer. However, for *memory* transfer, in which *synaptic weights* must be being transferred from one neural circuit (region) to another, what is the mechanism? Here, we present a synaptic transfer mechanism whose structure provides some understanding of the phenomena that have been implicated in memory transfer, including the nested oscillations at various frequencies. The circuit is based on the principle of pulse-gated, graded information transfer between neural populations.

PACS numbers: 87.18.Sn,87.19.lj,87.19.lm,87.19.lq

## INTRODUCTION

The case of Henry Molaison (H.M.) [1] taught us that there was a distinction between the location of short- and longterm memory storage in the brain. A bilateral medial temporal lobectomy removing the anterior part of H.M.’s hippocampi and other nearby brain regions resulted in an inability for H.M. to create new long-term memories (but he could still recall old memories), while leaving intact the ability to form and recall short-term memories. This caused researchers to believe that encoding and retrieval of longterm memories was mediated by distinct systems and that memories may be formed in one location in the brain, but consolidated elsewhere.

Sleep has been shown to support the consolidation of memory [2]. During non-rapid-eye-movement (NREM) sleep, thalamo-cortical spindles [3] and hippocampal sharp wave-ripples [4] have been implicated in declarative memory consolidation [5–11]. Evidence [8, 9, 12, 13] suggests that longterm memory consolidation is coordinated by the generation of hierarchically nested hippocampal ripples (100-250 Hz), thalamo-cortical spindles (7-15 Hz), and cortical slow oscillations (< 1 Hz) enabling memory transfer from the hippocampus to the cortex.

Consolidation has also been demonstrated in other brain tasks, such as in the acquisition of motor skills, where there is a shift from activity in prefrontal cortex to premotor, posterior parietal, and cerebellar structures [14] and in the transfer of conscious to unconscious tasks, where activity in initial unskilled tasks and activity in skilled performance are located in different regions, the so-called ‘scaffolding-storage’ framework [15].

Cortical oscillations are an important mechanism that enables information transfer in the brain. Experimental evidence has demonstrated improved communication between sender and receiver neurons when gamma-phase relationships exist between upstream and downstream populations [16–18]. Loss of theta-band coherence has been shown to result in memory deficits [19] and pharmacological enhancement can improve learning and memory [20]. Theoretical investigations support the idea that coincident spiking can allow neuronal populations to overcome their threshold and propagate spikes downstream [21–25].

Recent work shows that, by separating a neural circuit into a feedforward chain of gating populations and a second chain coupled to the gating chain (graded chain), graded information (i.e. information encoded in firing rate amplitudes) may be faithfully propagated and processed as it flows through the circuit [26–30]. The neural populations in the gating chain generate pulses, which push populations in the graded chain above threshold, thus allowing information to flow in the graded chain [29, 30].

The separation of a circuit into gating and graded chains, so-called ‘synfire-gated synfire chains’ (SGSCs), allows the flow of information through the graded chain component to be precisely controlled by the gating chain component. This control allows the circuit to process information [26, 27], make decisions [27], and, importantly, deploy learning in a precise manner [28].

Using SGSCs, circuits have been constructed that learn a covariance matrix from an autoregressive process by gating information through a circuit with a designated Hebbian learning module, enabling the encoding of the covariance matrix in a set of synapses [28].

In this paper, we will describe how a set of previously learned synapses may in turn be copied to another module with a pulse-gated transmission paradigm that operates internally to the circuit and is independent of the learning process.

## METHODS

### Model

We use a mean-field model of an SGSC,

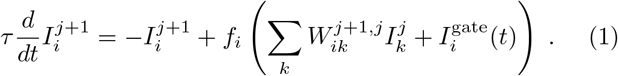

Here, 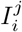 denotes the synaptic current in neural population *i* in layer *j*. 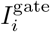 is a gating current, approximated by a square pulse of length *T*. Both currents are subthreshold. Thus, in the absence of the gating current, no information is propagated. For the simulations shown here, the time constant 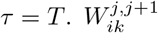 are elements of a synaptic connectivity matrix connecting layers *j* and *j* + 1. And *f* is a non-linear activity function, giving the firing rate, which, for an SGSC, is approximately piecewise-linear of the form

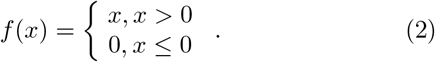

These equations may be derived from an underlying set of integrate-and-fire equations via a Fokker-Planck analysis [29].

Synaptic weights evolve according to a differential equation implementing Hebbian learning

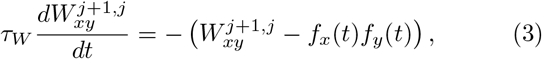

where *f_x_* and *f_y_* are firing rates of neuronal populations on post- and pre-synaptic (resp.) sides of a synapse, *W_xy_*. Here, *τ_W_* = 20,000 × 19*T*, where 19*T* is the length of a single instance of a repeated gating motif (see below).

Note carefully that firing rates in this system are turned on and off by gating current pulses, 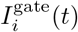. For Hebbian learning, synaptic modification (other than an overall leak) will only occur when firing occurs simultaneously on both sides of the synapse. Hence, learning may be turned on and off depending on the gating sequence used in the circuit. When information is gated to both pre- and post-synaptic sides of a synapse simultaneously, synaptic weights are modified. When information is gated *through* the synapse (i.e. first a pulse to the pre-synaptic population, then a pulse to the post-synaptic population, giving graded information propagation) no synaptic modification occurs.

The use of pulse-gating [28] is an alternative to other mechanisms, such as synaptic scaling, spike-timing dependent plasticity, and synaptic redistribution [31], that have been appealed to in order to regulate Hebbian learning, which is an inherently positive-feedback process, since effective synapses are strengthened, but ineffective synapses are weakened. The advantage of pulse-gating in this context is the capability of precisely controlling the destination and timing of information propagation, and hence learning, in the circuit.

### Connectivity and Pulse Sequence

The neural circuit connectivity used for synaptic copy is shown in Fig. 1. One sequence of a periodically repeating set of gating pulses for the control of information propagation through the circuit is shown in Fig. 2.

**FIG. 1.**
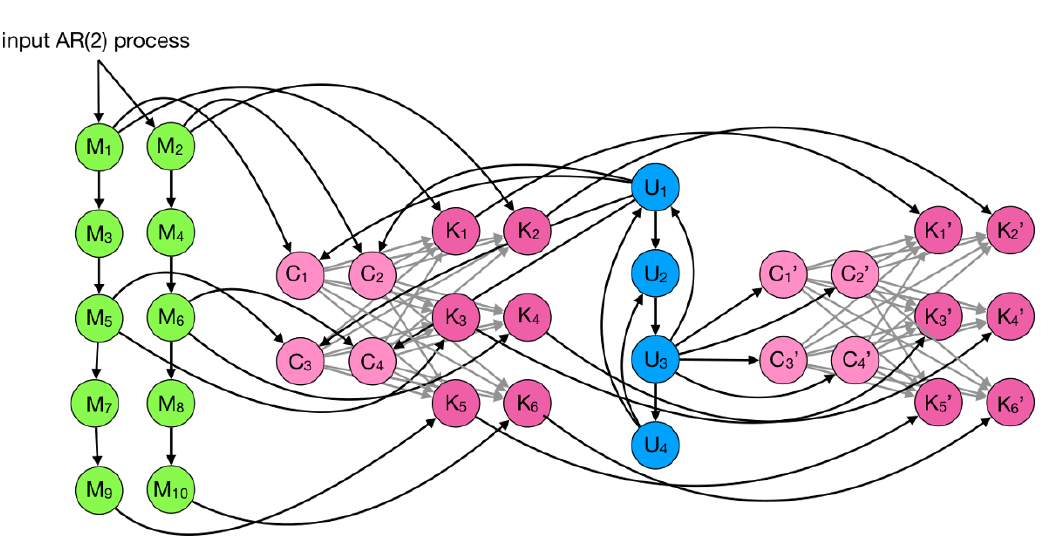
Synaptic copy circuit. *M*_1_ - *M*_10_ (green) denote neuronal populations used for a short-term memory. *M*_1_, *M*_3_, etc. denote populations that carry positive values of the input AR(2) process. *M*_2_, *M*_4_, etc. carry the absolute values of negative values of the input process (we use a zero mean process). *C* (pink) and *K* (magenta) populations represent populations pre- and post-synaptic to a set of Hebbian synapses. These synapses (light gray) are used to learn the lagged-covariances from the input process. *U* populations (blue) are used to coordinate a unit input value to various neural populations. *C*′ (pink) and *K*′ (magenta) populations are used for synaptic copy. The synaptic weights learned in the C and K populations are transferred to these populations. Synaptic weights for dark gray arrows are fixed. Arrows show synaptic direction.

**FIG. 2.**
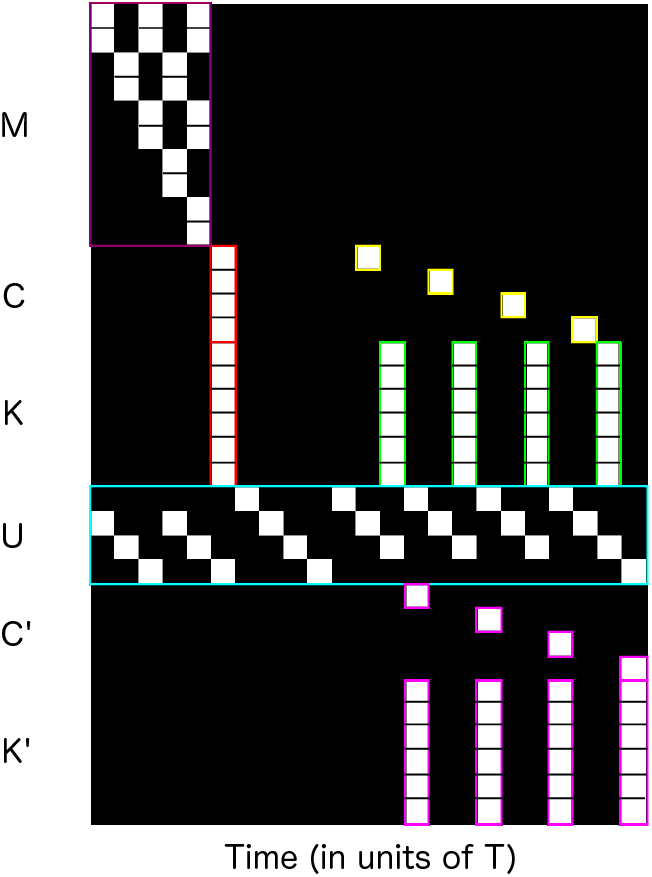
Synaptic copy pulse sequence. Each row corresponds to a gating pulse time series. Neuronal populations are denoted on the left: *M* (short-term memory), *C* (input to Hebbian synapses that learn AR(2) covariance), *K* (output from Hebbian synapses that learn AR(2) covariance), *U* (populations that coordinate unit inputs to various neural populations), *C*′ (input to Hebbian synapses to which AR(2) covariance is copied), and *K*′ (output from Hebbian synapses to which AR(2) covariance is copied). Purple boxes surround pulses that gate the AR(2) process through the short-term memory. Red boxes surround the pulses used to learn the AR(2) covariance. Yellow boxes gate Euclidean basis vectors to the *C* populations. Green boxes gate transformed Euclidean vectors to the *K* populations. The cyan box denotes pulses that gate a unit value through a subcircuit. Magenta boxes simultaneously gate Euclidean basis vectors and their transforms to *C*′ and *K*′, hence implementing synaptic copy.

In the first phase of operation of the circuit, a stochastic, autoregressive (AR(2)) process with covarying lags at times 2*T* and 4*T* is input to neural populations *M*_1_ and *M*_2_. Positive values (above the mean) are input to *M*_1_, and the absolute values of negative values are input to *M*_2_. Although the functionality is not explicitly used for these simulations, both positive and negative values are necessary to reconstruct an AR process from the covariance [28]. These values are propagated through short-term memories, *M*_3_ through *M*_10_.

The gating pulse sequence guides information through the circuit. Initial pulses propagate information from the input AR(2) process through the short-term memory populations *M*_1_ through *M*_10_ (Fig. 2, purple box at upper left). The neural populations *M*_3_, *M*_4_, and *M*_7_, *M*_8_ are temporary storage to allow neural currents to relax (and hence maintain accurate graded information propagation) between repeated information transfers to each population. Once the memory values are simultaneously in populations *M*_1_, *M*_2_, *M*_5_, *M*_6_, and *M*_9_, *M*_10_, the values are gated simultaneously to both sides, *C* and *K*, of a set of Hebbian synapses (Fig. 2, vertical red box).

Initially, no information is read into the *U* populations, and therefore, no further downstream processing occurs. During this phase of information processing, the covariance of the AR(2) process is encoded in the synapses between populations C and K of the neural circuit. This phase is meant to represent short-term memory encoding.

During the second phase of operation, the input process continues to be read into short-term memory. For this circuit, this is necessary in order to maintain the amplitude of the covariance encoded between the *C* and *K* populations. Otherwise, the leak would cause an exponential decrease in the covariance amplitude (although relative covariances would still have the correct proportional values). In a complete biological neural circuit, a means of stabilizing the synapses, such as lengthening the relaxation time constant, *τ_W_*, would need to be invoked.

The copy is begun with the input of a unit pulse into the *U*_1_ population. This pulse circulates through the *U* populations and is gated to the *C*_1_, *C*_2_, *C*_3_, and *C*_4_ populations successively (Fig. 2, yellow boxes). This corresponds to writing the Euclidean basis vectors, (1, 0, 0, 0), (0,1, 0, 0), etc., to the input of the covariance. The unit pulse is then gated through the *C-K* synapses (Fig. 2, green boxes) producing a transformed unit vector in the K populations. The unit pulse then gates the Euclidean basis vectors and the transformed vectors simultaneously to both sides of the synapses connecting the *C^r^* populations with the *K*′ populations. Upon multiple repeats of this process, Hebbian learning then encodes a copy of the C-K synapses in the *C*′-*K*′ synapses. This phase of operation is meant to represent consolidation of short-term memory to long-term memory.

## RESULTS

In Fig. 3, we show the AR(2) process values gated through the circuit for both encoding and copy phases. Fig. 3A shows input during the encoding phase as it is gated through the short-term memory populations, *M*, and simultaneously gated to *C* and *K* populations, hence encoding the covariance. Fig. 3B shows circuit operation after the copy phase has begun. At this point, the unit pulse may be seen circulating through the *U* populations, and the Euclidean basis vectors and their transforms being generated in the *C* and *K* populations, and their simultaneous propagation to *C*′ and *K*‡ populations leading to synaptic copy. Fig. 3C,D show sets of repeated encoding and copy sequences.

**FIG. 3.**
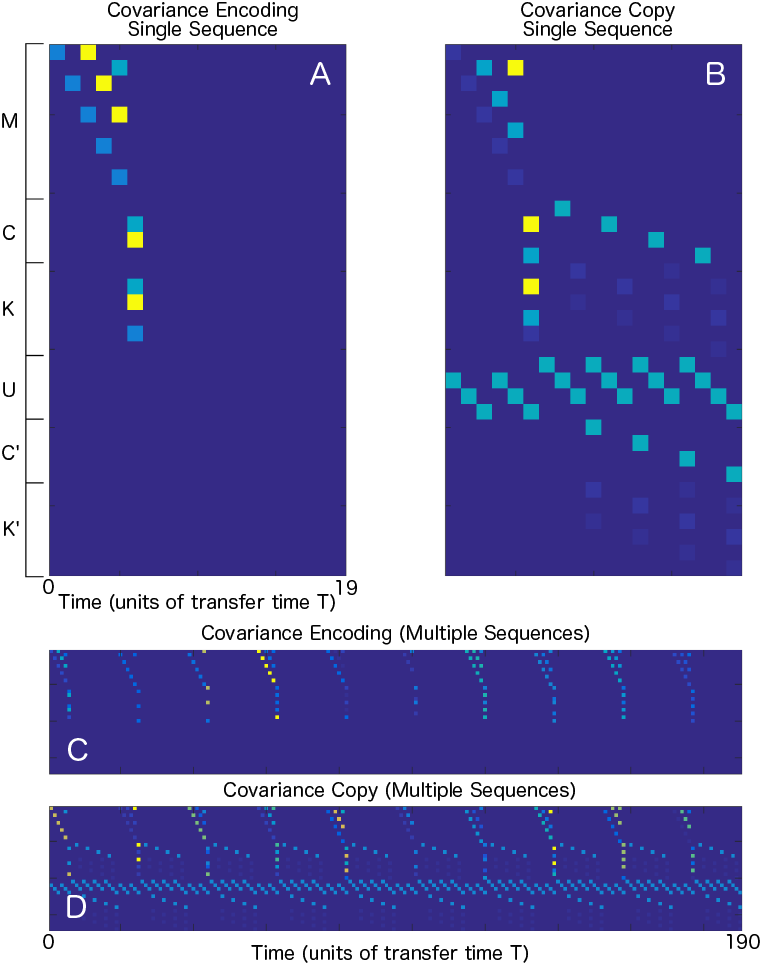
Synaptic copy graded sequences, encoding and copy phases. Each row corresponds to a graded pulse time series. Neuronal populations are denoted on the left: *M, C, K, U, C′*, and *K*^1^. A) Graded information pulses during the encoding phase. Here, the time series is read in and copied simultaneously to pre- and post-synaptic populations *C* and *K* (resp.). During this phase, the lagged covariance of the AR(2) time series is encoded via Hebbian learning in synapses between these populations. B) Graded information pulses during the copy phase. Here, the encoded covariance is maintained using the same pulses as in A). Additionally, a unit pulse is written into the *U* populations, and this unit pulse is used to write Euclidean basis vectors to the *C* populations, the unit vectors are gated through the covariance encoding synapses, and their transforms result in the *K* populations. The unit vectors and transforms are then written simultaneously to pre- and post-synaptic populations *C*′ and *K*′ (resp.). Hebbian learning causes the covariance to be copied to these synapses. Note that the graded pulses, of length T, here appear flat. This is because the simulation was long, and only the peak of the pulse was saved. See [26] for the canonical graded pulse shape.

**FIG. 4.**
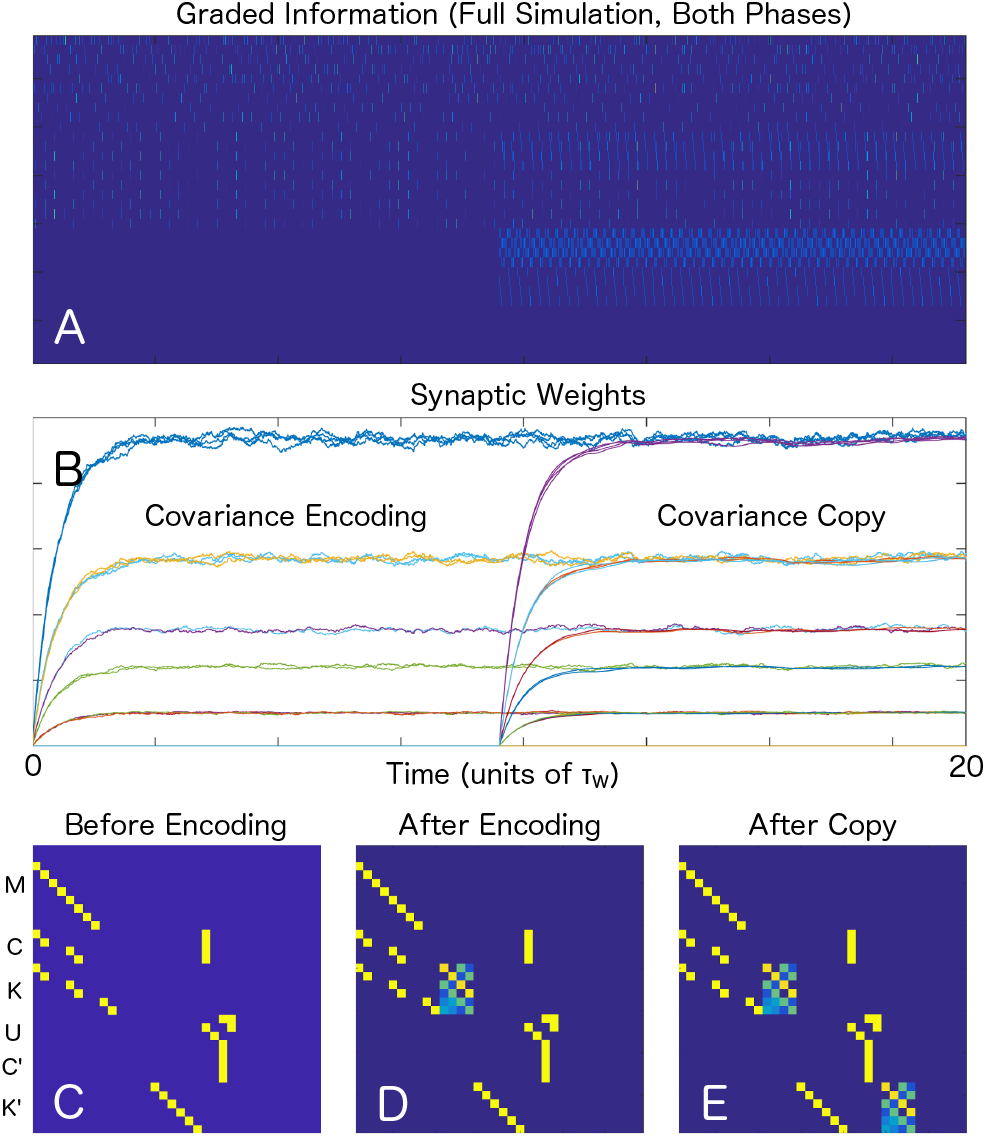
Synaptic weight evolution. A) Graded pulses shown for the full synaptic copy simulation. The encoding and copying phases are evident in the lower half of the plot, where the onset of the copy pulses may be seen. B) The synaptic weights shown as a function of time. The weights encoding the covariance from the stochastic time series input are seen transiently rising to their asymptotic values first. Then, once the unit pulse has been introduced and the copying phase has been initiated, the weights encoding the copied covariance are seen rising to their asymptotic values. Note that the copied weights rise smoothly because of the effective double filtering of the Hebbian learning evolution. First from the initial encoding, then a second smoothing from the copy. C, D, E) The synaptic weights shown as they evolve within the connectivity matrix: before encoding, after encoding, after copy. Note that the copied synapses both in panel E) and in panel B) have been normalized by an overall multiplicative scalar that is induced by the differing effective leakages from the periodic learning sequences (times between simultaneous copy to pre- and post-synaptic populations).

In Fig. 4A, we show the graded information as it is repeatedly propagated through the circuit 20 × *τ_W_* times. Halfway through the simulation, at time 10 × *τ_W_*, the synaptic copy phase is turned on. In Fig. 4B, we see the evolution of the encoding and copy synapses, with the encoding phase occurring first, and the copy phase initiating after the encoded synapses are stable. In Fig. 4C, D, E we show the synaptic connectivities (see Fig. 1) with the covariance matrices initially, after encoding, and after copy (resp.).

## DISCUSSION

The synaptic copy circuit that we have presented here has an important feature that is remarkably similar to that seen experimentally in measurements of memory consolidation [8, 9, 12, 13], *viz.* nested, in-phase oscillations at multiple frequencies. During memory consolidation, that is, during the copy phase of our simulations, intermediate frequency oscillations are used to generate transformed vectors from encoding synapses that are propagated to copy synapses. High frequency oscillations are used to coordinate the transformation and propagation of information from one part of the circuit to another. These are all nested within the base frequency set by the overall sequence execution period of 19*T*.

We do not mean to imply that there is a direct mapping between our circuit and memory consolidation circuits in the brain. Our circuit is simply meant as an existence proof that synaptic copy is possible using the synfire-gated synfire chain mechanism in a neural circuit with local, Hebbian learning. In the circuit presented here, the size of the synaptic matrix being copied determines the intermediate frequencies responsible for generating the transformed Euclidean vectors in the copy process and also influences the low-frequency behavior. Thus, the relative nested oscillation frequencies involved in the copy vary depending on specifics of what is being copied. Nonetheless, the similarities in oscillatory epiphenomena are intriguing and the underlying concept, that the nature of the algorithm informs the structure of oscillatory phenomena seen in the circuit, could provide a way forward for reverse engineering information processing features of neural circuits from the details of spectral measurements.

An important feature of this synaptic copy mechanism is that it functions independently of how the information being copied was learned originally. This is important in that, if there were such a dependence, for instance, in the mammalian brain, then one would not expect a canonical set of frequencies to be involved in consolidation during NREM sleep. Indeed, the frequency content would vary depending on the day-to-day processes that the encoding system was exposed to. In the case considered here, one would need to re-generate the AR process whose covariance was learned initially. This would be impractical, particularly for a general purpose system, such as the brain obviously has.

In addition to being a useful model for understanding how memory is transferred between brain areas, pulse-gating and the capability of synaptic copy within a neural circuit may be useful in neuromorphic systems for which learning or communication is expensive, or for neuromorphic implementation of machine learning structures such as neural Turing machines [32]. For such circuits, it may be cheaper to temporarily enable (possibly expensive) learning only in specific regions of a neuromorphic chip (or chips). This would allow synapses to be copied to regions that, for instance, would be used for inference, but not intensive learning, similarly to how TPUs are used to make inferences that were learned on a GPU.

Finally, the synaptic copy circuit is an explicit solution to the long-standing ‘weight transfer problem’ [33, 34] that has hindered the implementation of biophysically realistic deep neural networks [35].

## AUTHOR CONTRIBUTIONS

Y.S. and B.W. contributed equally to this paper.

## Acknowledgements

L.T. thanks the Los Alamos National Laboratory (LANL) and the UC Davis Mathematics Department for their hospitality. This work was supported by the Natural Science Foundation of China grants 31771147 (Y.S., B.W., L.T.), 91232715 (L.T), by the Open Research Fund of the State Key Laboratory of Cognitive Neuroscience and Learning grant CN-LZD1404 (Y.S., B.W., L.T.), by the Beijing Municipal Science and Technology Commission under contract Z151100000915070 (Y.S., B.W., L.T.), and by the ASC Beyond Moore’s Law project at LANL (A.T.S.). LANL preprint - LA-UR-18-24103.

